# Inferring evolutionary relationships among *Crenotia* species (Bacillariophyta): Evidence from natural populations and monoclonal strains from Slovakia

**DOI:** 10.64898/2026.04.13.718240

**Authors:** Alica Hindáková, Pavla Urbánková, Jana Kulichová

## Abstract

Diatoms exhibit remarkable diversity in valve morphology, with the raphe system being a fundamental feature in classification of raphid pennate diatoms. The repeated loss of one of the two raphes during evolution has led to multiple independent origins of monoraphid diatoms. The phylogenetic affinities of the monoraphid genus *Crenotia* A. Z. Wojtal, erected from *Achnanthidium thermale* Rabenhorst, have not yet been clarified with molecular data. In this study, natural populations of *Crenotia* and monoclonal strains derived from them were examined using morphological observations and multilocus phylogenetic analyses based on nuclear and plastidial molecular markers. Three species of the genus *Crenotia* form a well-supported clade placed within a subgroup of monoraphid genera, which are closely related to Cymbellales D.G. Mann and other biraphid diatoms. This study establishes the first molecular framework for representatives of the genus *Crenotia*, demonstrating their monophyly and congruent interspecific relationships recovered with multiple molecular markers. The low intraspecific sequence variability and substantial interspecific divergence, together with clear morphological and ecological differentiation, support the recognition of the three investigated species.

## INTRODUCTION

The classification of diatoms has traditionally relied on comparative morphology. One of the defining characteristics of the order Achnanthales Silva is the presence of a raphe on one valve and the absence of a raphe on the second valve of the frustule (Round *et al*. 1990). This morphological feature is associated with the term heterovalvar frustules or monoraphid diatoms (Round *et al*. 1990). Recent molecular and cladistic studies have shown that the loss of one of the two raphes has occurred multiple times independently during diatom evolution (Kociolek & Stoermer 1986; Kulikovskiy *et al*. 2016; Nakov *et al*. 2018; Kulikovskiy *et al*. 2019; Alverson *et al*. 2025). The shared evolutionary history of monoraphids with biraphid diatoms, together with the polyphyletic nature of some monoraphid taxa, has further blurred traditional taxonomic boundaries (see e.g., Cox 2006; Cox & Williams 2006; Davidovich *et al*. 2016; Thomas *et al*. 2016; Ashworth *et al*. 2017; Kulikovskiy *et al*. 2020; Gorecka *et al*. 2021; Riaux-Gobin *et al*. 2021).

The complexity of monoraphid systematics highlights the need for targeted phylogenetic studies at the genus level, especially for taxa whose classification is defined solely by morphological criteria. An example of one such genus is *Crenotia* A.Z. Wojtal, which was established in 2013 based on morphological re-examination of the type material of *Achnanthidium thermale* Rabenhorst. Consistent with earlier observations by Lange-Bertalot & Ruppel (1980), Wojtal (2013) recognised two key diagnostic features: a zig-zag arrangement of areolae forming the striae and the bending of the frustule apparent when viewed from the girdle (Figs 1–3). The genus name *Crenotia* reflects the spring-dwelling (crenophilous) nature of these diatoms. Early records of *Crenotia* taxa (formerly assigned to *Achnanthidium* Kützing or *Achnanthes* Bory) come from mineral and thermal springs or freshwater and saline habitats worldwide (e.g., Rabenhorst 1864; Hustedt 1930; Krasske 1925, 1927; Poretzky & Anisimova 1933; Hustedt 1949; Lange-Bertalot & Krammer 1989). In recent decades, *Crenotia* species have been increasingly reported from mineral springs, gravel pit lakes, and streams throughout Europe (Wojtal 2013; Lai *et al*. 2019; Beauger *et al*. 2023) and Asia (Liu *et al*. 2020; Na *et al*. 2024). These records, including those compiled in AlgaeBase (Guiry & Guiry 2026), identify *Crenotia* as a diverse, globally distributed genus. The aims of our study were to describe the morphology of both natural populations and monoclonal cultures, to determine the phylogenetic placement of *Crenotia* representatives, and to compare traditional species boundaries with molecular data. To achieve these objectives, we collected benthic diatom communities from two types of habitats in Slovakia – mineral springs and a gravel pit lake, where populations of different *Crenotia* species, including *C. thermalis* (Rabenhorst) Wojtal, had previously been recorded (Hindáková 2009; Hindák & Hindáková 2013, 2014, 2015).

**Figure 1.**
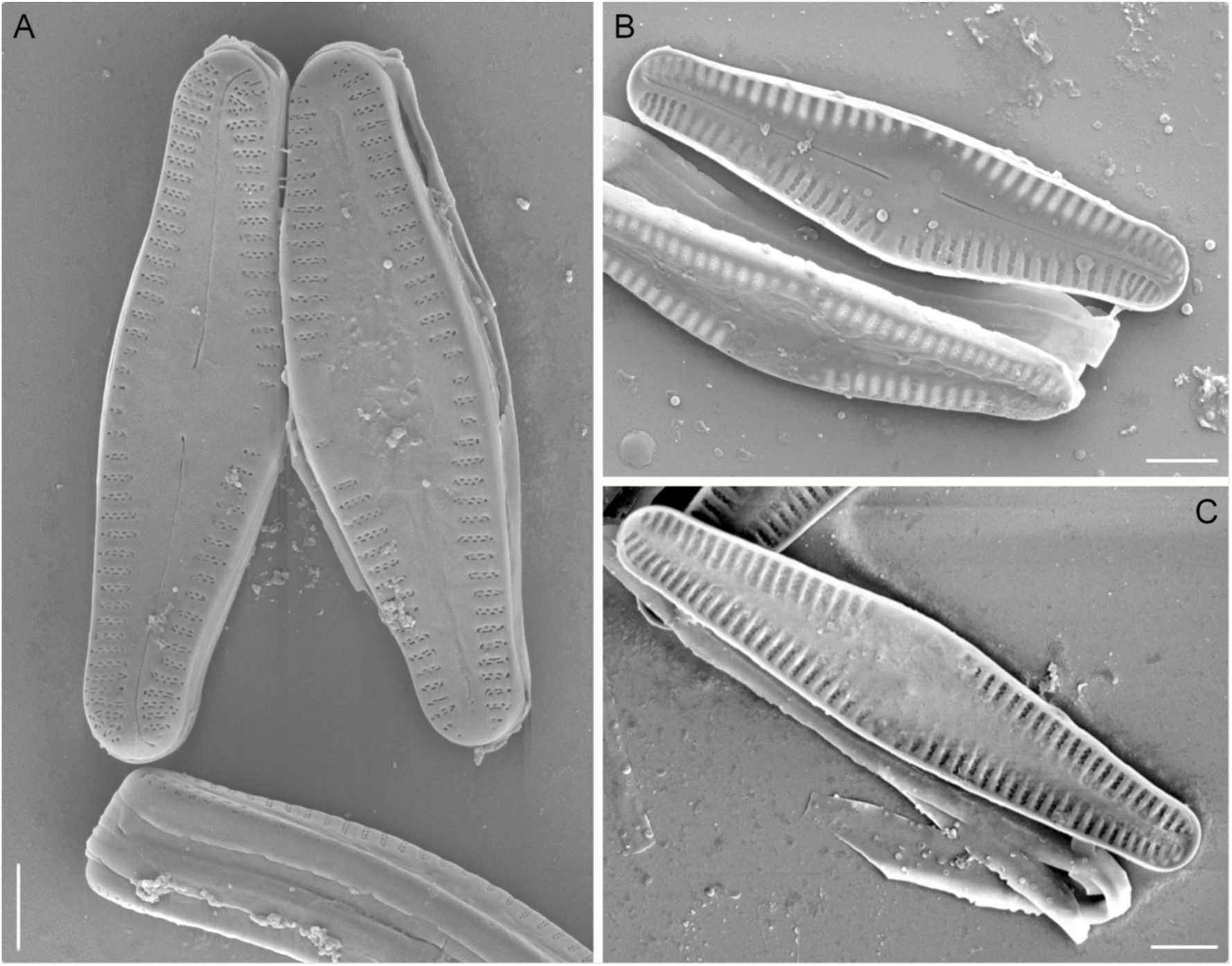
SEM microphotographs of *C. thermalis*. **A**, external view, raphe and rapheless valve, strain UH4-04D; **B**, internal view, raphe valve, strain UH4-04D; **C**, internal view, rapheless valve, strain UH2-02E. For strain details see Table S1. Scale bars: 2 µm.

**Figure 2.**
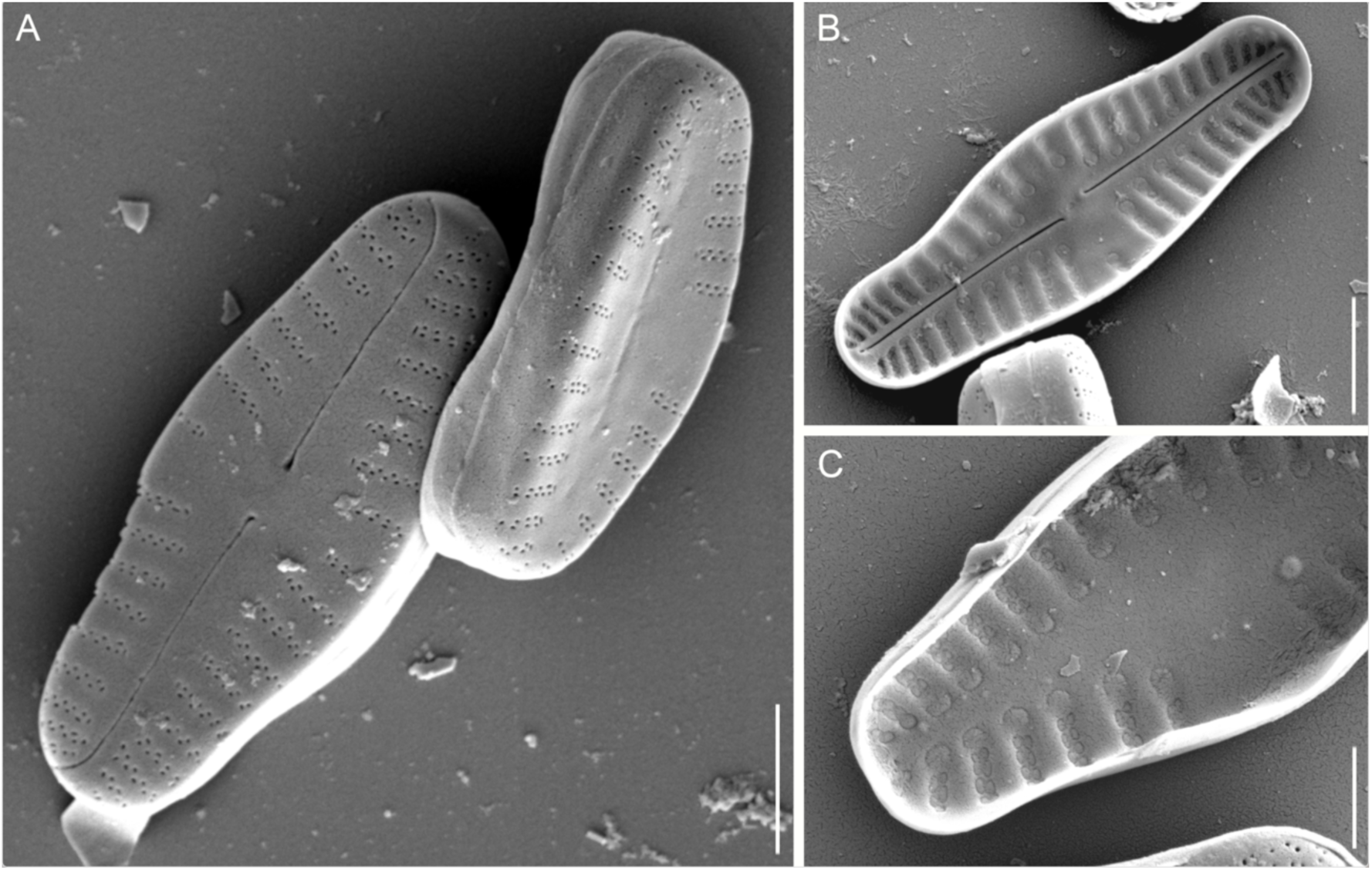
SEM microphotographs of *Crenotia* sp. **A**, external view, raphe and rapheless valve, strain UH9-06B; **B**, internal view, raphe valve, strain UH9-06B; **C**: internal view, rapheless valve, strain UH9-06B. For strain details see Table S1. A, B, scale bars: 2 µm, C, scale bar: 1 µm.

**Figure 3.**
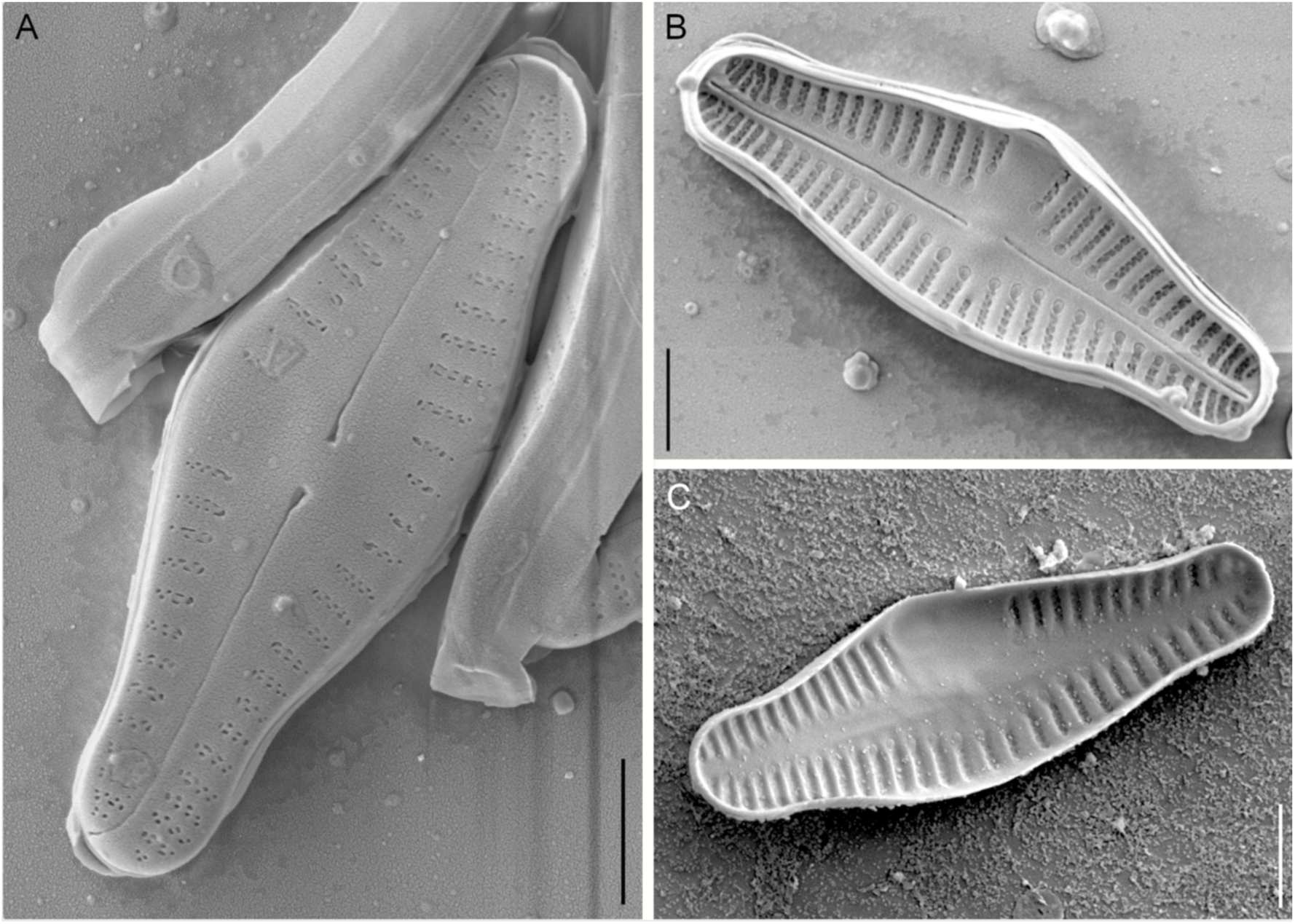
SEM microphotographs of *C. rumrichorum*. **A**, external view, raphe valve, strain UH7-03F; **B**, internal view, raphe valve, strain UH7-03F; **C**, internal view, rapheless valve, strain UH7-02F. For strain details see Table S1. Scale bars: 2 µm.

## MATERIAL AND METHODS

### Collection of the material

Benthic diatom communities were collected from five locations (Fig. S1) in the period 2021–2022 (Table 1). At Močiar (#1) – an active geothermal borehole on a travertine fen, diatoms formed macroscopically visible colonies in shallow travertine cascades and on moist travertine crusts. At locality Rojkov (#2) – an active spring crater known as Zlá Voda (’The Bad Water’), benthic diatoms were abundant along the natural banks of the travertine pool. At an active geothermal borehole on travertine bedrock behind the village Gánovce (#3), the shallow pool and upper channels were covered by mats of filamentous cyanobacteria among which diatoms were abundant. At Sivá Brada (#4) – an active natural geyser on a travertine pile, fine brownish layers of diatoms covered the surface of the travertine crusts. And at Zlaté Piesky (#5) – a gravel pit lake in the urban area of the city of Bratislava, brown diatom mats along the lake shore were macroscopically visible during cold late winter days.

**Table 1.** Measurements of physico-chemical parameters of water at sampling sites.

| **Locality** | **#1 Močiar** | **#2 Rojkov** | **#3 Gánovce** | **#4 Sivá Brada** | **#5 Zlaté piesky** |
| --- | --- | --- | --- | --- | --- |
| **Sampling date** | July 2021 | July 2021 | September 2022 | September 2022 | March 2022 |
| **GPS - North** | 49.1540358 | 49.1467436 | 49.0272508 | 49.0072500 | 48.1850739 |
| **GPS - East** | 19.1517761 | 19.1604350 | 20.3350783 | 20.7232611 | 17.1869439 |
| **Altitude (m a.s.l.)** | 442 | 445 | 620 | 487 | 130 |
| **Sampling site** | main borehole | crater | main borehole | main geyser | gravel pit lake |
| **Temperature (°C)** | 18.1 | 21.9 | 21.8 | 14.6 | 6.1 |
| **pH** | 6.3 | 6.5 | 6.6 | 6.5 | 8.8 |
| **Cond.<sup>1</sup> (mS/cm)** | 3.3 | 4.2 | 3.6 | 5.6 | 0.84 |
| **TDS<sup>2</sup> (g/l)** | 1.77 | 2.1 | 1.79 | 2.3 | 0.42 |
| **Resistivity (Ω)** | 280 | 238 | 279 | 180 | 1170 |
| **Salinity (ppt)** | 1.78 | 2.14 | 1.82 | 2.95 | 0.38 |

<sup>1</sup>Cond: Conductivity, <sup>2</sup>TDS: Total Dissolved Solids.

The water from all four springs (#1–4) was slightly acidic (Table 1) and strongly mineralised, with a sulphate–hydrocarbonate composition and high calcium concentrations (Dítě *et al*. 2011). Locations 1, 3, and 4 (Močiar, Gánovce, and Sivá Brada), with temperatures ranging from 14.7 °C to 21 °C, are classified as thermal springs due to temperatures higher than the average annual air temperature in the region (Meinzer 1923), whereas location 2 (Rojkov) does not have a narrow annual temperature range (own measurements; data not shown), despite similar hydrochemistry. The water of the gravel pit lake (#5, Zlaté Piesky) represents a different type of habitat, with much lower mineralisation and higher pH (Table 1). Groundwater from the Danube river, which is filtered through sand and gravel bedrock, originally had a slightly alkaline oligotrophic character, which changed to mesotrophic due to human recreational activities (Hindáková 2009). The water parameters were measured in the field using the portable pH tester Hanna HI 98107 and the conductivity meter Hanna HI 98192.

The collected fresh material was transported to the laboratory in plastic containers. One part was used to isolate monoclonal cultures, another was fixed for long-term storage, and the rest was used to prepare permanent slides according to the standard hydrogen peroxide method (Taylor *et al*. 2007). Light microscopic (LM) examination and photo documentation of natural populations of the genus *Crenotia* was carried out at the Institute of Botany, Plant Science and Biodiversity Centre, Slovak Academy of Sciences (PSBC, SAS) using a Leitz Diaplan light microscope with Wild Photoautomat MPS45. The documentary and voucher materials of the field samples are stored at the PSBC, SAS.

### Isolation of monoclonal strains

Monoclonal strains of the *Crenotia* species were isolated by single-cell picking using an ultrathin Pasteur pipette. The selected cell was washed through a series of drops of sterile WC medium (Guillard & Lorenzen 1972) to minimize the risk of potential contamination. After several days of cultivation in 96-well tissue chambers in circumneutral WC medium at 18° C under cool-white permanent fluorescent lights, part of the biomass of uncontaminated isolates was harvested for LM and DNA isolation.

Another part of the biomass was inoculated into glass tubes and maintained under long-term cultivation conditions (8° C, light shading with filter paper). The voucher material was oxidized directly on the coverslips according to Trobajo & Mann (2019).

Coverslips were mounted on slides with Naphrax (Brunel Microscopes, Wiltshire, UK) and photographed with an Olympus BX51 microscope with differential interference contrast (DIC), using Olympus Z5060 digital microphotographic equipment (Olympus Corporation, Tokyo, Japan). Selected strains, oxidised on coverslips, were mounted onto metal stubs, coated with platinum, and examined under a JEOL IT800 high resolution scanning electron microscope (SEM). The voucher material, living monoclonal strains, and frozen biomass are stored at the Phycology Research Group, Department of Botany, Charles University.

### Amplification of specific DNA regions

DNA was isolated from 10 µL of biomass using 20 µL of InstaGene^TM^ Matrix Buffer (Bio-Rad Laboratories, Hercules, CA, USA) following the manufacturer’s instructions. The Polymerase Chain Reaction (PCR) using MyTaq^TM^ Hot Start master mix containing 0.2 µL MyTaq polymerase, 4.0 µL of 5× MyTaq reaction buffer, and 0.25 µL each of forward and reverse primers (25 pmol/µL); template DNA (1–5 µL) and nuclease-free water were added to a final volume of 20 µL. Different primer combinations were used to amplify nuclear and plastidial DNA regions: 18S rDNA - 34F (Thüs *et al*. 2011) and ITS5-DR (Urbánková & Veselá 2013), D1-D2 of 28S rDNA - LSU-80DF and LSU-740DR (Veselá *et al*. 2012), *rbc*L - DPrbcL1 and DPrbcL7 (Daugbjerg & Andersen 1997), *psb*A - psbA-F and psbA-R1 (Yoon *et al*. 2002). The PCR conditions for 18S amplification were as follows: initial denaturation at 94° C - 5 min; 35 cycles of 94° C - 1 min, 51° C - 1 min, 72° C - 2 min; final extension at 72° C - 10 min. For 28S: initial denaturation at 95° C - 1 min; 35 cycles of 95° C - 20 s, 51° C - 30 s, 72° C - 30 s; final extension at 72° C - 5 min. For *rbc*L: initial denaturation at 95° C - 1 min; 35 cycles of 95° C - 20 s, 51° C - 30 s, 72 ° C - 55 s; final extension at 72° C for 10 min. For *psb*A: initial denaturation at 94° C - 1 min; 35 cycles of 94 °C - 30 s, 54° C - 30 s, 72° C - 30 s; final extension at 72° C - 7 min. PCR products were purified by SPRI-select Bead for Size Selection (Beckman Coulter, Brea, CA, USA) following the manufacturer’s protocol. Sequencing was performed by Macrogen Europe B.V. (Amsterdam, Netherlands). Of the four loci analysed for *Crenotia*, *rbc*L was selected as the primary marker due to its highest phylogenetic resolution. 28S rDNA and *psb*A were used as secondary markers because they showed slightly lower but still comparable variability. The highly conserved 18S rDNA was selectively sequenced, as it was not expected to provide additional resolution once lineages had been distinguished using the more variable markers.

### Phylogenetic reconstructions

The raw sequences of twenty *Crenotia* strains were visualised and edited in Geneious Prime ver. 2022.2.2 (www.geneious.com), retaining only high-quality reads for phylogenetic analyses. All sequences were deposited in the Barcode of Life Data Systems (BOLD; strain IDs: CRE001-26–CRE020-26), and a subset of long reads was additionally deposited in the National Center for Biotechnology Information (GenBank accession numbers: 18S rDNA: PZ163566–PZ163570; 28S rDNA: PZ163571–PZ163574; *rbc*L: PZ143698–PZ143704; *psb*A: PZ143680–PZ143683) (Table S1). To determine the phylogenetic position of *Crenotia*, we compiled a dataset of our newly obtained unique sequences and GenBank sequences (18S rDNA, 28S rDNA, *rbc*L, *psb*A, *psb*C) from raphid pennate diatoms, with Eunotiaceae Kützing used as the outgroup (Table S2). Protein-coding genes (*psb*A, *psb*C, *rbc*L) were aligned using the Geneious aligner. The 18S rDNA sequences were processed using SSU-Align ver. 0.1.1 (Nawrocki 2009) with the default parameters and the eukaryotic secondary structure model, which aligns conserved regions and removes poorly aligned positions based on the reference secondary structure. The 28S rDNA sequences were aligned analogously using R-Coffee (Wilm *et al*. 2008) with default parameters, which identifies conserved and highly variable regions based on secondary structure information. Poorly aligned regions in the 28S rDNA alignment were subsequently removed using TCS (Chang *et al*. 2015) with default settings. Concatenator ver. 0.2.1 (Vences *et al*. 2022) was used to concatenate markers and define partitions. The dataset was divided into five partitions: one for each rDNA gene (n=2) and one per codon position for the combined plastid genes (n=3). The concatenated alignment (216 taxa, 5748 base pairs) is available in the supplementary material.

The maximum likelihood phylogeny was calculated using the IQtree ver. 3.0.1 (Trifinopoulos *et al*. 2016) and a ModelFinder (Kalyaanamoorthy *et al*. 2017) to select the best-fitting substitution models under a partitioned scheme (option -m MFP+MERGE). Partitioned schemes and selected models are in Table S3. Node support was assessed with 1000 ultrafast bootstrap (UFB) replicates (option -bb 1000). Unlike traditional bootstrap, UFB provides more unbiased support values across a large number of gene trees. In this framework, a UFB ≥ 95% can be interpreted as strong support, which is roughly equivalent to a 95% probability that the clade is correct (Minh *et al*. 2013). Unstable taxa were evaluated using the R (R Core Team 2021) package ‘Rogue’ (Smith 2022), applying variants with one, two, and three taxa removed, but no rogue taxa were detected. The resulting tree was visualized and edited using iTOL (Letunic & Bork 2024). For clarity, only the subtree relevant to the phylogenetic relationships of the genus *Crenotia* and among the investigated species is shown (Fig. 7); the complete tree is provided in Figure S2.

Due to low support for the deep branches in the phylogeny, an Approximately Unbiased (AU) test (Shimodaira, 2002) implemented in IQ-TREE ver. 3.0.1 with 10,000 multiscale bootstrap replicates was used to test the relationship of *Crenotia* to other species (or groups of species) within a subgroup of monoraphid genera, which are closely related to Cymbellales and other biraphid diatoms. The seven tested topological constraints are listed in Table S4.

## RESULTS

### Morphological Characterisation

Natural populations and monoclonal strains derived from them share the morphological characteristics typical of the genus *Crenotia*, as outlined by Wojtal (2013). Our observations in SEM (Figs 1-3) and LM (Figs 4-6) provided additional details on their frustules, contributing to a more precise characterization of this genus.

**Figure 4.**
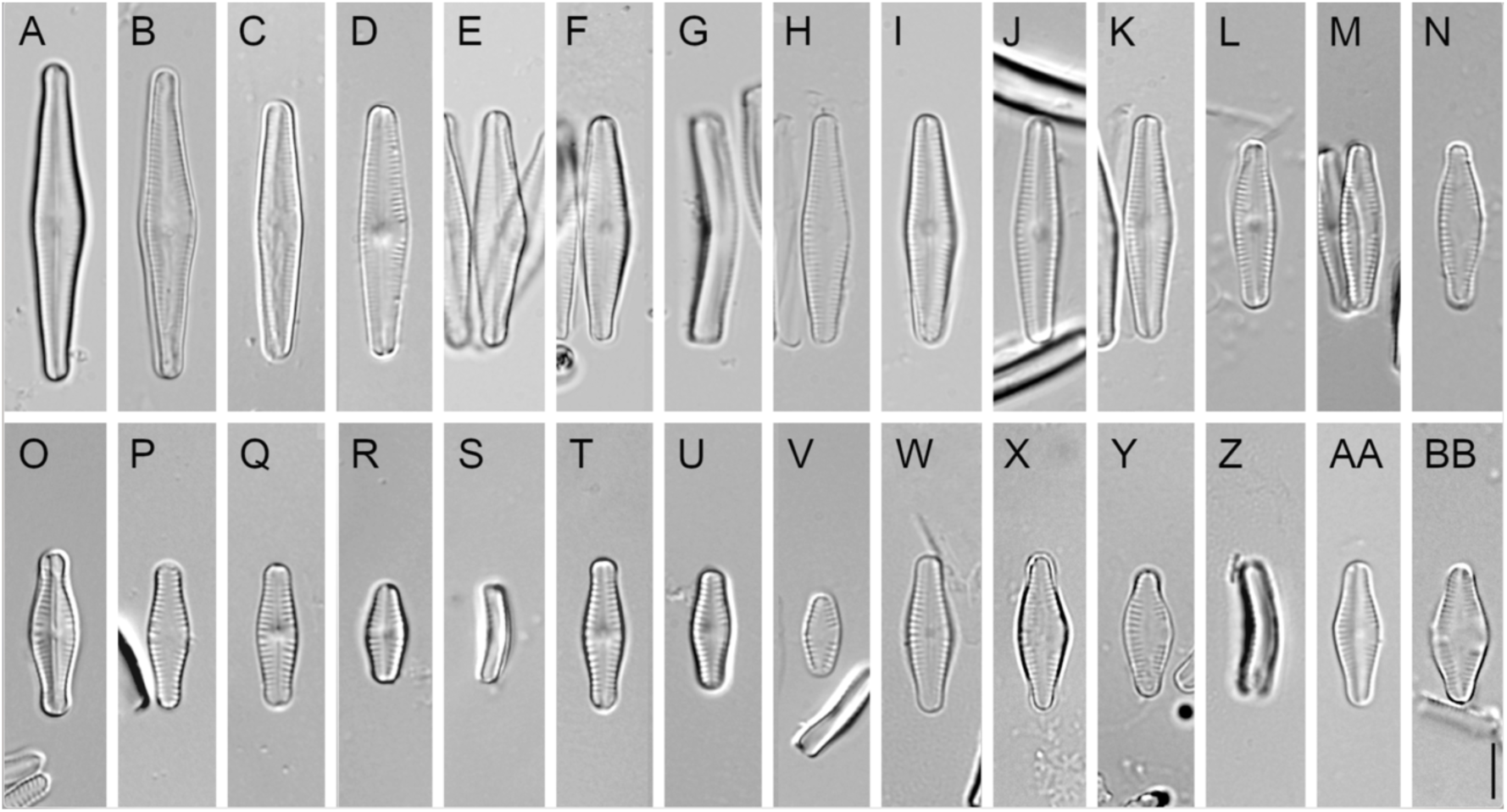
LM microphotographs of *C. thermalis* (A–N), *Crenotia* sp. (O–V), and *C. rumrichorum* (W–BB) strains. **A**, **B**, *C. thermalis*, strain UH4-07C; **C**, **D**, *C. thermalis*, strain UH4-03D; **E–G**, *C. thermalis*, strain UH4-07D; **H–K**, *C. thermalis*, strain UH4-02E; **L–N**, *C. thermalis*, strain UH4-04D; **O–S**, *Crenotia* sp., strain UH9-06G; **T–V**, *Crenotia* sp., strain UH9-06B; **W–Z**, *C. rumrichorum*, strain UH7-02F; **AA, BB**, *C. rumrichorum,* strain UH7-03E. For strain details see Table S1. Scale bar: 5 µm.

Emended description. Frustules exhibited a species-specific degree of bending along the median plane, producing a saddle-shaped appearance and characteristic double image in LM (Figs 4G, S, Z; 5C, F, I). In the valve view, cells appeared slightly dorsiventral due to longitudinal asymmetry. Smaller, late-stage cells of the life cycle tended to be more elliptic in shape. The bevelled appearance of the valve ends, clearly visible in LM, results from the specific arrangement of the terminal striae. SEM revealed a flat lanceolate axial area on the raphe valve, with a poorly differentiated central area often extending to the valve margin (with at least 1–2 missing striae; Figs 1–3). The raphe was slightly undulate and eccentric, bending toward the side with curved terminal fissures. The rapheless valve had a broader lanceolate axial area and featured a distinct, wrinkled central depression extending toward the apices, visible as shadow-like markings in LM and forming a slightly convex internal band interrupted at the center in SEM. Striae on the raphe valve were uniseriate to biseriate, moderately radiate to parallel in the center, becoming more radiate and denser toward the apices.

**Figure 5.**
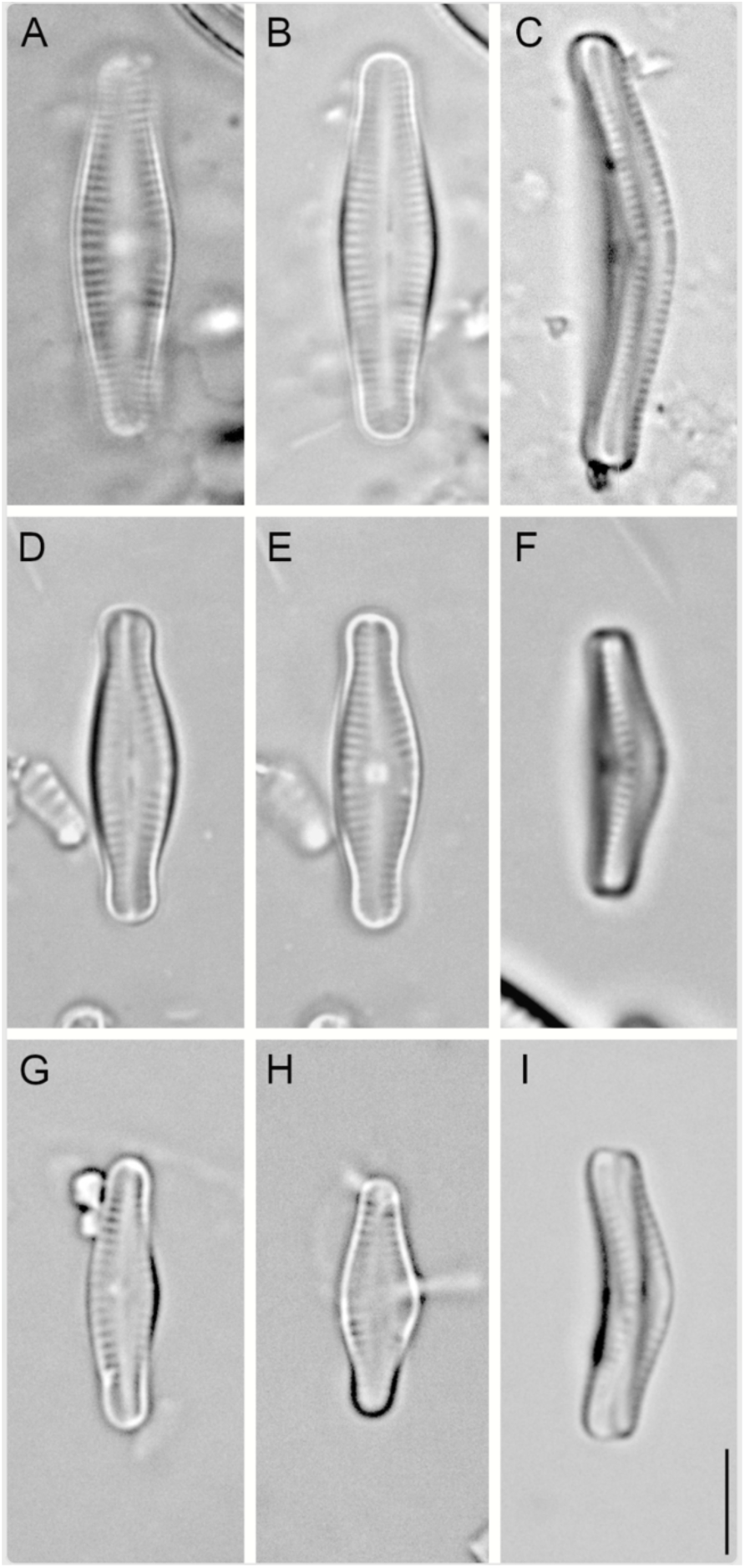
LM microphotographs of *Crenotia* populations from the field. **A–C**, *C. thermalis*, Močiar (#1); **D–F**, *Crenotia* sp., Sivá Brada (#4); **G–I**, *C. rumrichorum*, Zlaté piesky (#5). Scale bar: 5 µm.

These terminal striae were offset beneath the curved fissures, creating an asymmetrical pattern at the valve ends. A comparable asymmetry was present on the rapheless valve (notably in *C. hedinii* Rioual, Ector & Wetzel; Rioual *et al*. 2019), further contributing to the bevelled appearance. Transapical striae were short and varied in length, with terminal striae up to 0.3 µm longer, as seen in SEM (Figs 1–3). On the rapheless valve, striae were consistently biseriate, radiate to parallel, and evenly spaced.

Three species of the genus *Crenotia* were distinguished based on comparison of our material with observations reported in recent studies (Hindáková 2009, Wojtal 2013, Hindák & Hindáková 2013, 2014, 2015). *C. rumrichorum* (Lange-Bertalot) Wojtal (Figs 3; 5W–BB) exhibits a rimmed depression, *Crenotia* sp. (Figs 2; 5O–V) has strongly bent frustules in girdle view, and *C. thermalis* (Figs 1; 5A–N), is larger than the other two species and lacks the capitate and/or subrostrate apices. The variability in size and shape in *C. thermalis* initially made it unclear whether these differences reflected life-cycle allometry or interspecific divergence (Fig. 4A–N), but sequencing data supported an interpretation of intraspecific allometry. *Crenotia* sp. closely resembles *Achnanthes grimmei* var. *hyalina* Hustedt, as illustrated in Simonsen (1987, Plate 537: 6–11; LM). However, this variety has not been revised using modern taxonomic criteria, and it was never formally transferred to the genus *Crenotia*, which was established by Wojtal (2013). The original habitat of this taxon is likely ecologically quite different from the Slovak sampling site (#4 Sivá Brada, Table 1) as *A*. *grimmei* var. *hyalina* was frequently associated with stoneworts of the genus *Chara* Linnaeus and occasionally in a streamlet near an oasis on the Sinai Peninsula (Hustedt 1949). Due to the lack of type material examination and limited documentation, we refrain from assigning a definitive name to this species at this stage.

### Crenotia thermalis

Natural populations of *C. thermalis* were observed free-living or colonial, either attached to substrates or forming ribbon-like chains (Fig. S1C, F, I). The frustules were slightly bent along the transapical axis, with the raphe valve concave and the rapheless valve convex (Fig. 1A, B). Valves were lanceolate, lanceolate-linear, or elliptic-lanceolate in shape (Fig. 4A–N), measuring 9.8–34.7 µm in length and 3.0–5.2 µm in width. Valve ends were broadly rounded and only subtly subcapitate. The axial area was variably lanceolate, while the central area was not distinctly differentiated (Fig. 1A). It appeared symmetric or asymmetric, with unilateral expansion reaching the valve margin; in such cases, 1–2 striae were absent on the raphe valve and 4–5 on the rapheless valve (Fig. 1). Striae were biseriate, arranged parallel to slightly radiate in the central part of the valve, with a density of (19)–20–22(–23) in 10 µm, becoming more radiate and denser toward the apices (Fig. 1). On the raphe valve, the last 5–7 striae near the apices were arranged most densely (33–40 in 10 µm).

Populations of *Crenotia thermalis* from Močiar (Fig. 5A–C), Rojkov, and Gánovce (#1–3) exhibited moderate variability in valve outline, with frustules from Rojkov (#2) showing the closest resemblance to the type material of *Achnanthidium thermale* Rabenh. published in Wojtal (2013), Plate 28: 1–17, 29: 1–7, 30: 1, 2. Compared to the natural material, the 14 monoclonal strains examined were within the same size range (14.0–27.9 µm / 3.5–5.0 µm) and tended to form a gibbous centre. Ultrastructural features corresponded to natural populations. In the living cell, the brownish chloroplast bearing one pyrenoid lies adjacent to the raphe-less valve, sometimes extending below the raphe-valve (Fig. 6A–E). It occupies almost the whole cell, and its margins are slightly to markedly lobed. There are two or more oil droplets in the cell.

**Figure 6.**
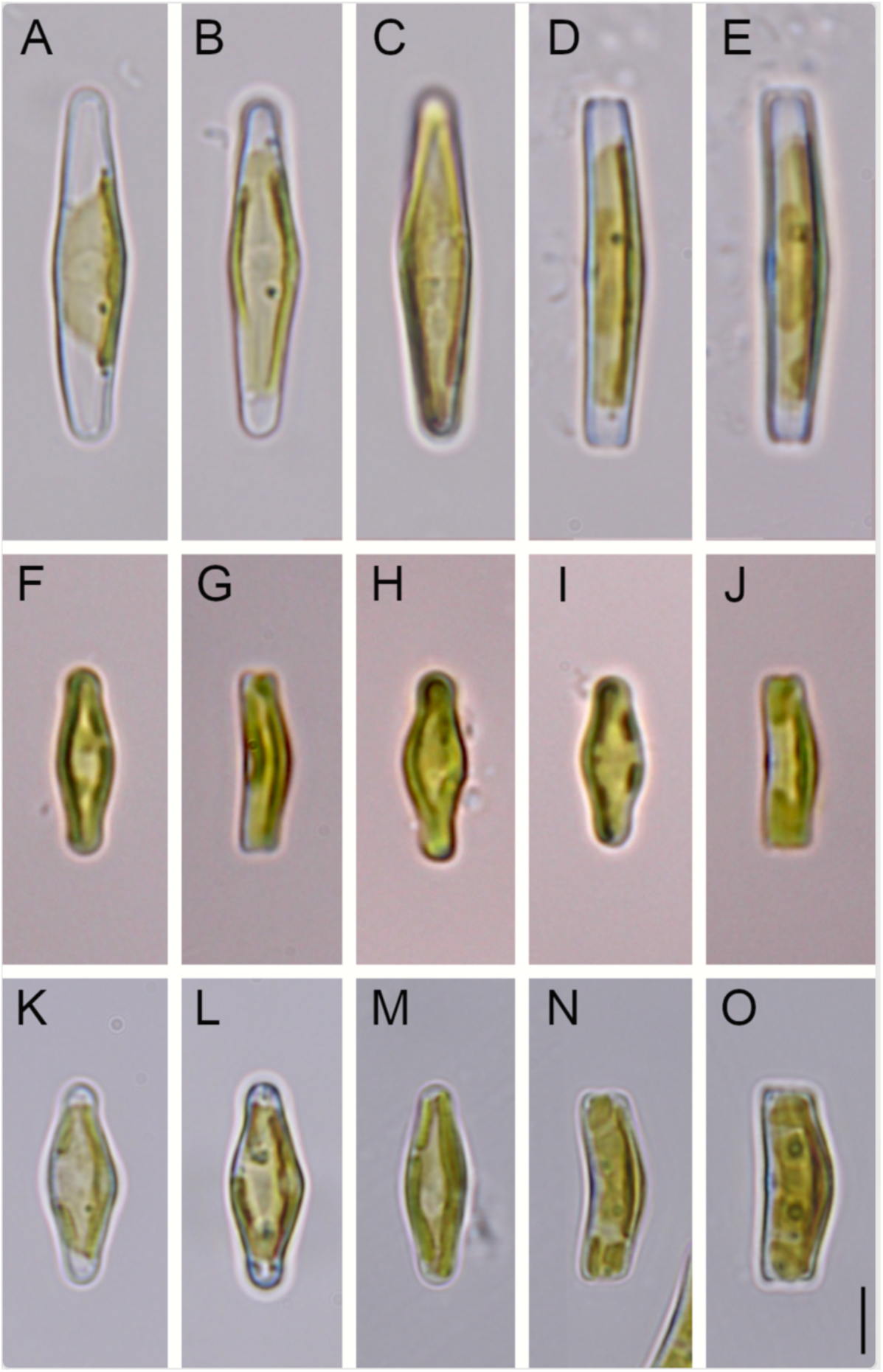
LM Microphotographs of living *C. thermalis* (A–E), *Crenotia* sp. (F–J), and *C. rumrichorum* (K–O) strains. **A–E**, *C. thermalis*, strain UH4-07C; **F**, **G**, *Crenotia* sp., strain UH9-06B; **H–J**, *Crenotia* sp., strain UH9-06G; **K**, **L**, *C. rumrichorum*, strain UH7-03F; **M–O**, *C. rumrichorum*, strain UH7-2F. For strain details see Table S1. Scale bar: 5 µm.

### Crenotia sp

Cells of *Crenotia* sp. were either free-living or attached to substrates (Fig. S1L). The species shares with *C. thermalis* strongly bent frustules, lanceolate to elliptic-lanceolate outlines, and indistinct central areas that can be symmetrically or asymmetrically expanded (Fig. 2). Other characteristics of *Crenotia* sp. were different. The number of ‘missing’ striae in the expanded central area was 1–2 in both valves (Fig. 2). It had small valves (8.3–14.9 µm long, 3.4–4.3 µm wide) and capitate to subrostrate valve ends (Fig. 4O–V). Striae were predominantly biseriate, occasionally uniseriate in the central region, radiate, and counted 18–22(–23) in 10 µm (Fig. 2). On the raphe valve, the last 3–5 striae toward the apex were the densest.

Compared to the natural material from Sivá Brada (#4), the three monoclonal strains exhibited a broader size range (7.1–15.2 µm / 2.9–4.4 µm), were less strongly bent along the transapical axis, often showed a slightly gibbous central area, and occasionally formed ribbon-like colonies. The chloroplast (Fig. 6F–J) looked very similar to that of *C*. *thermalis*.

### Crenotia rumrichorum

Natural populations of *C. rumrichorum* were solitary or in pairs, either free-living or attached to the substrate by a mucilage stalk secreted from the distal end of the raphe (Fig. S1O). Similarly to *Crenotia* sp., *C. rumrichorum* had strongly bent frustules, lanceolate to elliptic-lanceolate outlines, and indistinct central areas that could be symmetrically or asymmetrically expanded (Fig. 3). The number of ‘missing’ striae in the central area of *C. rumrichorum* was 2–3 on the raphe valve and 5–6 striae on the rapheless valve. It had small valves (8.3–18.5 µm long, 3.1–4.6 µm wide), often swollen in the center, giving them a gibbous and slightly asymmetric appearance, and terminated in broadly subrostrate ends (Fig. 4W–BB). Striae of *C. rumrichorum* were biseriate or occasionally uniseriate in the center, short, slightly parallel, and number 20–29 in 10 µm, with the last 4–6 striae on the raphe valve being the densest (Fig. 3). Internally, the rapheless valve consistently exhibited a rimmed depression in place of the ‘missing’ striae (Fig. 3C), which appeared as a shadow under LM (Fig. 4Y, BB).

Population from Zlaté Piesky (#5) corresponded to *Achnanthes thermalis* var. *rumrichorum* Lange-Bertalot (Lange-Bertalot & Krammer 1989, Plates 77:30–38; 78:2–4; LM and SEM), now regarded as a synonym of *Crenotia rumrichorum*. In the three monoclonal strains examined, frustules were slightly broader (11.5–15.3 µm / 3.4–5.7 µm), the rimmed depression on the rapheless valve was less distinct, and striae density was lower (20–22 in 10 µm). The chloroplast (Fig. 6K–O) was very similar to those of the previous two species.

### Phylogenetic Relationships and Genetic Differences

Because monoraphid diatoms originated multiple times independently within biraphid lineages (e.g., Alverson *et al*. 2025), the phylogenetic position of *Crenotia* was evaluated using a dataset including representatives of major monoraphid and biraphid groups. The concatenated alignment contained a substantial proportion of missing data; however, exploratory analyses indicated that their inclusion did not affect the overall topology. In this phylogeny, *Crenotia* species formed a well-supported lineage (UFB = 100) within a clade containing Cymbellales D.G. Mann, Lyrellales D.G. Mann, *Rhoicosphenia* Grunow and other biraphid taxa (Fig. 7). Despite the robust support for the sister position of *Crenotia* to *Planothidium* Round & Bukhtiyarova ex Round (UFB = 98) and the highest log-L for this topology in the AU test, alternative placements of *Crenotia* to other closely related monoraphid genera cannot be excluded (Table S4). The only topology that produced significantly worse results was one in which *Crenotia* was a sister lineage to the *Achnanthidium minutissimum* species complex (Fig. 7).

**Figure 7.**
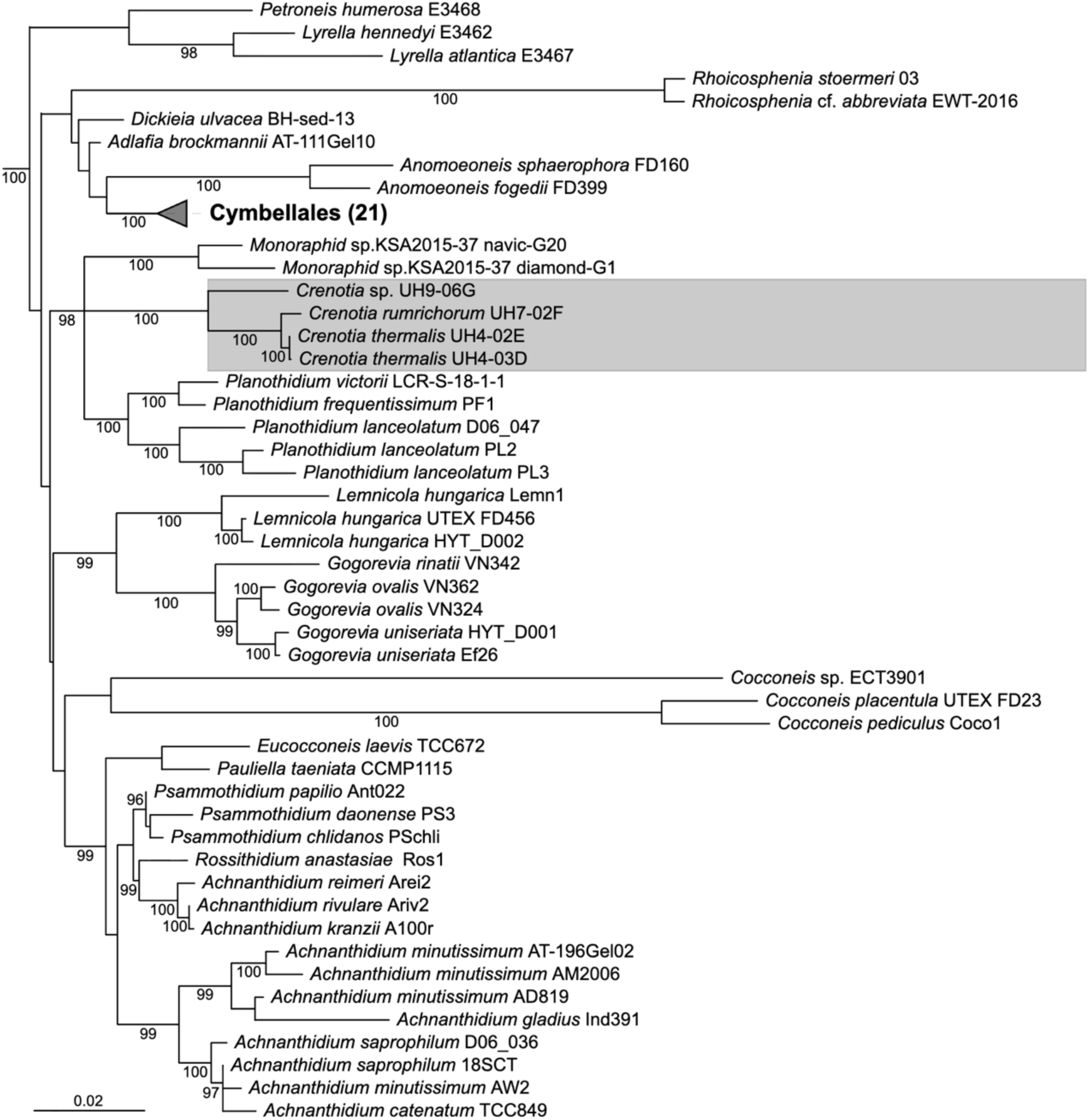
Maximum likelihood (ML) phylogeny showing the placement of the genus *Crenotia* (grey box) among its monoraphid relatives and related biraphid taxa. The tree is based on a concatenated dataset of nuclear (18S rDNA, 28S rDNA) and plastid (*rbc*L, *psb*A, *psb*C) markers. Only the subtree relevant to the placement of *Crenotia* is shown; the complete phylogeny is provided in Figure S2. Statistical support values (UFB > 95; 1000 ultrafast bootstrap replicates) are indicated at the nodes.

Analysis of molecular data obtained from twenty strains confirmed the existence of three distinct species within the genus *Crenotia* (Fig. 7). No intraspecific variability was observed in the molecular markers across the three lineages, with one exception. In the *C*. *thermalis* lineage, the *rbc*L marker differed by a single base pair in a sequence longer than a thousand base pairs. The observed difference in the *rbc*L marker likely reflects population-level genetic variation rather than distinct species. The two *rbc*L variants within *C. thermalis* were not distinguishable based on morphology (Figs 1; 4A, B, H–K, L–N vs. Fig. 4C, D, E–G) and occurred in sympatry (Table S1).

### Ecological preferences based on natural samples

The differentiation of the three *Crenotia* species was also reflected in their environmental preferences. *C*. *thermalis* occurred in travertine mineral springs with moderately high ion concentrations (#1-3) and *Crenotia* sp. was found in a similar spring environment but with much higher ion concentrations (#4). *C*. *rumrichorum* inhabited a gravel pit lake (#5), which had distinctly different hydrochemical conditions, including higher pH, greater temperature fluctuations and lower ion concentrations (Table 1).

## DISCUSSION

This study combines detailed morphological observations with multilocus phylogenetic analyses to clarify species boundaries and evolutionary relationships among representatives of the diatom genus *Crenotia*. Our observations using light and scanning electron microscopy have confirmed that all specimens conform to the generic concept of *Crenotia* as defined by Wojtal (2013), thus allowing a more precise circumscription of the genus. A comprehensive morphological analysis was conducted on the key frustule features, including bending along the median plane, valve shape, striae arrangement, and ultrastructural characteristics of raphe-bearing and rapheless valves. The field material and derived monoclonal strains were found to group consistently into three species: *C. thermalis*, *C. rumrichorum*, and *Crenotia* sp. The latter is morphologically similar to *Achnanthes grimmei* var. *hyalina*, but a definitive identification requires a thorough re-examination of Hustedt’s original material. While all species exhibit distinctive traits, such as biseriate zig-zag striae, a rimmed depression on rapheless valves was observed in *C. rumrichorum*, a feature also reported in other monoraphid genera.

Historically, several *Crenotia* taxa have been included in *Achnanthidium* or *Achnanthes*, as evidenced by their synonymy (Guiry & Guiry 2026). This is understandable, as *Achnanthes* has been widely used as a generic name for monoraphid diatoms, and the classification of *Achnanthidium* and *Achnanthes* with early taxonomic concepts was based solely on observations of living cells, distinguishing free-living (*Achnanthidium*) from stalked forms (*Achnanthes*; Kützing 1844; Rabenhorst 1853). Nevertheless, our data showed that natural populations of *Crenotia* may form ribbon-like colonies or be solitary – either free-living or attached to the substrate via mucilage stalks (Fig. S1), particularly when growing among densely packed cyanobacterial *Rivularia* C. Agardh filaments in recent stromatolites (Hindák & Hindáková 2013). Although the examined *Crenotia* species has a similar valve shape and frustules bent along the median plane as the *Achnanthidium minutissimum* species complex, the AU test ruled out their close relationship. Our phylogeny suggests that the genus *Planothidium* is the closest relative of the genus *Crenotia*; however, an alternative placement of the three *Crenotia* species with other monoraphid genera - within a clade containing Cymbellales - was similarly plausible (Table S4). A notable common feature between these genera is the marginal rimmed depression present in *C. rumrichorum*, the closest known relative of *C*. *thermalis,* and in some *Planothidium* species (*P*. *reichardtii* Lange-Bertalot & Werum, *P*. *lanceolatum* (Brébisson ex Kützing) Lange-Bertalot, and *P*. *amphibium* C.E.Wetzel, Ector & L.Pfister). However, other *Planothidium* species, including those with resolved phylogenies (Jahn *et al*. 2017), differ in the structure of the central area of the rapheless valves. Furthermore, marginal rimmed depression is not only found in *Planothidium* and *Crenotia*, but also occurs in representatives of other monoraphid genera, e.g., *Platessa strelnikovae* Enache, Potapova & Morales (Enache *et al*. 2014) and *Psammothidium lauenburgianum* (Hustedt) Bukhtkiyarova & Round (Bukhktiyarova & Round 1996). The unresolved phylogenetic relationships within the part of the tree relevant to the placement of *Crenotia* emphasise the need for expanded molecular and morphological datasets. Nevertheless, the monophyly of *Crenotia* was consistently and well supported across all analyses, and relationships among the three investigated species were stable. Within *C. thermalis*, LSU and *psb*A did not resolve the single-base-pair *rbc*L variant, indicating low intraspecific divergence. Despite the relatively high proportion of missing data in the concatenated alignment, exploratory analyses indicated that the placement of *Crenotia* was stable. Limited support for deeper nodes is consistent with a transcriptome-based study showing that this part of the diatom phylogeny remains poorly resolved (Alverson *et al*. 2025).

Interpreting ecological preferences and distribution of *Crenotia* species is complicated by historical shifts in diatom species concept (Mann 1999, 2010). Taxa assigned to *Achnanthidium thermalis / Achnanthes thermale* underwent several taxonomic revisions, in which morphologically similar taxa were synonymised without considering potential habitat differences or the remoteness of their places of origin (Lange-Bertalot & Ruppel 1980; Lange-Bertalot & Krammer 1989; Krammer & Lange-Bertalot 1991; Lange-Bertalot *et al*. 1996). Limited or absent microphotographic documentation in many historical records makes it difficult to verify taxonomic identity. The reported broad geographical and environmental ranges of *A. thermalis / A. thermale* (Guiry & Guiry 2026) are therefore likely to reflect lumped taxa that are predominantly found in thermal and/or mineral springs. More recent studies employing both LM and SEM have demonstrated that some *Crenotia* species can inhabit geographically restricted but ecologically diverse habitats, such as high-altitude streams, hot springs, and standing waters in Tibet (Liu *et al*. 2020; Na *et al*. 2024; Rioual *et al*. 2019). Our results corroborate the ecological breadth of *Crenotia* species and demonstrate that phylogenetic relationships do not necessarily reflect habitat similarity. *C. rumrichorum*, the closest relative of *C. thermalis*, inhabits a man-made lake with low mineralisation and alkaline pH, whereas *Crenotia* sp., despite occurring in habitats similar to those of *C. thermalis*, forms a sister lineage to these species. The frequent occurrence of *Crenotia* species in thermal and/or mineral springs indicates that this relatively rarely reported genus may be a valuable indicator of ecologically specific environments.

## ACKNOWLEDGEMENTS

AH extends her thanks to the Slovak Scientific Grant Agency VEGA (no. 2/0064/25) for its support. PU is co-financed from the state budget by the Technology agency of the Czech Republic within the SIGMA Programme (TQ03000438). JK is supported by the Charles University Research Program (Cooperatio) and acknowledges the Viničná Microscopy Core Facility (VMCF of the Faculty of Science, Charles University), an institution supported by the MEYS CR (LM2023050 Czech-BioImaging).

---

**Table S1**. List of *Crenotia* strains represented by at least one molecular marker. GenBank accession numbers correspond to long-read sequences deposited in the National Center for Biotechnology Information. BOLD IDs refer to strain records in the Barcode of Life Data Systems, where associated sequences are available.

**Table S2**. GenBank accession numbers of sequences included in the phylogenetic analyses. Column A-B: name of the taxon in Figures (Figs 7; S2) and in the alignment, respectively. Columns C-G: molecular markers and their position in the alignment (bp).

**Table S3**. Partitioning scheme of the concatenated alignment and selected substitution models. Position of plastidial markers (*rbc*L, *psb*C, *psb*A) is indicated in Table S2.

**Table S4**. AU test assessing the position of *Crenotia* within a subgroup of monoraphid genera, which are closely related to Cymbellales and other biraphid diatoms (Fig. 7). LogL = log-likelihood of data given the un-/constrained topology, p-value = AU test probability (values <0,05 that indicate rejection are in bold), RSS = bootstrap approximation diagnostic (lower values are more reliable), Constraint = clade constraint used to generate the topology (groups are explained in Table S2 and Fig. 4).

**Figure S1**. Photographs of localities, sampling sites, and phytobenthic communities. **A–C**, Močiar (#1); **D–F**, Rojkov (#2); **G–I**, Gánovce (#3), **J–L**, Sivá Brada (#4); **M–O**, Zlaté Piesky (#5). Scale bar: 20 µm.

**Figure S2**. Maximum likelihood (ML) phylogeny of the pennate raphid diatoms calculated with the IQtree program (Trifinopoulos et al. 2016). The ML phylogeny is based on a concatenated data set of the nuclear-encoded regions (18S rDNA, 28S rDNA) and the plastid genes (*rbc*L, *psb*A, and *psb*C). Statistical support UFB >95 (1000 UltraFast bootstrap replicates) is indicated at the nodes of the tree. Newly obtained *Crenotia* sequences are in a grey box.

**Alignment**. The concatenated sequence alignment (216 taxa, 5748 base pairs) of five molecular markers. The positions are as follows: 1–1524: *rbc*L; 1525-2748: *psb*C; 2749–3702: *psb*A; 3703–4171: 28S rDNA; 4172–5748: 18S rDNA. Details regarding taxon names and their GenBank accession numbers are provided in Table S2.

## REFERENCES

Alverson A. J., Roberts W. R., Ruck E. C., Nakov T., Ashworth M. P., Bryłka K., Downey K.M., Kociolek J.P., Parks M., Pinseel E., Theriot E. C., Tye S. P., Witkowski A., Beaulieu J.M. & Wickett N.J. 2025. — Phylogenomics reveals the slow-burning fuse of diatom evolution. Proceedings of the National Academy of Sciences of the United States of America 122 (22): e2500153122. 10.1073/pnas.2500153122

Ashworth M. P., Lobban C. S., Witkowski A., Theriot E. C., Sabir M. J., Baeshen M. N., Hajarah N. H., Baeshen N. A., Sabir J. S. & Jansen R. K. 2017. — Molecular and morphological investigations of the stauros-bearing, raphid pennate diatoms (Bacillariophyceae): *Craspedostauros* Ej Cox, and *Staurotropis* Tbb Paddock, and their relationship to the rest of the Mastogloiales. Protist 168 (1): 48–70. 10.1016/j.protis.2016.11.001

Beauger A., Voldoire O., Allain E., Gosseaume P., Blavignac C., Baker L.-A. & Wetzel C. E. 2023. — Biodiversity and environmental factors structuring diatom assemblages of mineral saline springs in the French Massif Central. Diversity 15 (2): 283. 10.3390/d15020283

Bukhtiyarova L. & Round F. E. 1996. — Revision of the genus *Achnanthes sensu lato*. Psammothidium, a new genus based on A. marginulatum. Diatom Research 11 (1): 1–30. 10.1080/0269249X.1996.9705361

Cox E. J. 2006. — *Achnanthes sensu stricto* belongs with genera of the Mastogloiales rather than with other monoraphid diatoms. European Journal of Phycology 41(1): 67–81. 10.1080/09670260500491543

Cox E. J. & Williams D. M. 2006. — Systematics of naviculoid diatoms (Bacillariophyta): a preliminary analysis of protoplast and frustule characters for family and order level classification. Systematics and biodiversity 4 (4): 385–399. 10.1017/S1477200006001940

Daugbjerg N. & Andersen R. A. 1997. — A molecular phylogeny of the heterokont algae based on analyses of chloroplast-encoded *rbc*l sequence data. Journal of Phycology 33 (6): 1031–1041. 10.1111/j.0022-3646.1997.01031.x

Davidovich N. A., Davidovich O. I., Witkowski A., Li C., Dabek P., Mann D. G., Zgłobicka I., Kurzydłowski K. J., Gusev E. & Górecka E. 2016. — Sexual reproduction in *Schizostauron* (Bacillariophyta) and a preliminary phylogeny of the genus. Phycologia 56 (1): 77–93. 10.2216/16-29.1

Dítě D., Dražil T. & Janák M. 2011. — Management Model for the Carpathian Travertine Salt Meadows. [Manažmentový model pre Karpatské travertínové slaniská]. Daphne, Institute of Applied Ecology., Bratislava, 16 p.

Enache M. D., Potapova M. & Morales E. A. 2014. — *Platessa strelnikovae* (Bacillariophyta), a new species from Maine and Vermont lakes, USA. Beihefte zur Nova Hedwigia 143: 239–244.

Górecka E., Ashworth M. P., Davidovich N., Davidovich O., Dąbek P., Sabir J. S. & Witkowski A. 2021. — Multigene phylogenetic data place monoraphid diatoms *Schizostauron* and *Astartiella* along with other fistula-bearing genera in the Stauroneidaceae. Journal of Phycology 57 (5): 1472–1491. 10.1111/jpy.13192

Guillard R. R. L. & Lorenzen C. J. 1972. — Yellow-green algae with chlorophyllide C. Journal of Phycology 8: 10–14. 10.1111/j.1529-8817.1972.tb03995.x

Guiry M. & Guiry G. 2026. — AlgaeBase. World-wide electronic publication: http://www.algaebase.org

Hindák F. & Hindáková A. 2013. — Mass development of phototrophic microorganisms near a thermal geyser at Gánovce. Limnologický spravodajca 7 (1): 11–16.

Hindák F. & Hindáková A. 2014. — Cyanobacteria and algae of mineral springs on a travertine pile of Sivá Brada (Spiš/Zips, Eastern Slovakia). Limnologický spravodajca 8 (2): 27–33.

Hindák F. & Hindáková A. 2015. — Cyanobacteria and algae of mineral springs of the fen Močiar at Stankovany, Central Slovakia. Bulletin Slovenskej botanickej spoločnosti 37 (2): 161–197.

Hindáková A. 2009. — On the occurrence of *Achnanthes thermalis* var. *rumrichorum* (Bacillariophyceae) in Slovakia. Fottea 9: 193–198. 10.5507/fot.2009.020

Hustedt F. 1930. — Bacillariophyta (Diatomeae). Zweite Auflage. Verlag von Gustav Fischer, Jena, 466 p. (Die Süsswasser-Flora Mitteleuropas)

Hustedt F. 1949. — Diatomeen von Sinai-Halbinsel und aus dem Libanon-Gebiet. Hydrobiologia 2: 24–55.

Chang J.-M., Di Tommaso P., Lefort V., Gascuel O. & Notredame C. 2015. — TCS: a web server for multiple sequence alignment evaluation and phylogenetic reconstruction. Nucleic Acids Research 43 (W1): W3–W6. 10.1093/nar/gkv310

Jahn R., Abarca N., Gemeinholzer B., Mora D., Skibbe O., Kulikovskiy M., Gusev E., Kusber W.-H. & Zimmermann J. 2017. — *Planothidium lanceolatum* and *Planothidium frequentissimum* reinvestigated with molecular methods and morphology: four new species and the taxonomic importance of the sinus and cavum. Diatom Research 32 (1): 75–107. 10.1080/0269249X.2017.1312548

Kalyaanamoorthy S., Minh B. Q., Wong T. K. F. von Haeseler A. & Jermiin L. S. 2017. — ModelFinder: fast model selection for accurate phylogenetic estimates. Nature Methods 14: 587–589. 10.1038/nmeth.4285

Kociolek, J. P. & Stoermer, E. F. 1986. — Phylogenetic relationships and classification of monoraphid diatoms based on phenetic and cladistic methodologies. Phycologia 25 (3): 297–303. 10.2216/i0031-8884-25-3-297.1

Krammer K. & Lange-Bertalot H. 1991. — Bacillariophyceae: Achnanthaceae. Kritische Ergänzungen zu *Navicula* (Lineolatae) und *Gomphonema*. Gustav Fischer Verlag, Jena, 437 p. (Süßwasserflora von Mitteleuropa; 2/4).

Krasske G. 1925. — Die Bacillariaceen-Vegetation Niederhessens. Abhandlungen und Berichte des Vereins für Naturkunde zu Kassel, 7-79.

Krasske G. 1927. — Diatomeen deutscher Solquellen und Graderwerke. Archiv für Hydrobiologie 18: 252–272.

Kulikovskiy M. S., Andreeva S. A., Gusev E. S., Kuznetsova I. V. & Annenkova N. V. 2016. — Molecular phylogeny of monoraphid diatoms and raphe significance in evolution and taxonomy. Biology Bulletin 43(5): 398–407.

Kulikovskiy M., Maltsev Y., Andreeva S., Glushchenko A., Gusev E., Podunay Y., Ludwig T.V., Tusset E. & Kociolek J. P. 2019. — Description of a new diatom genus *Dorofeyukea* gen. nov. with remarks on phylogeny of the family Stauroneidaceae. Journal of Phycology 55(1): 173–185. 10.1111/jpy.12810

Kulikovskiy M., Maltsev Y., Glushchenko A., Kuznetsova I., Kapustin D., Gusev E., Lange-Bertalot H., Genkal S. & Kociolek J. P. 2020. — *Gogorevia*, a new monoraphid diatom genus for *Achnanthes exigua* and allied Taxa (Achnanthidiaceae) described on the basis of an integrated molecular and morphological approach. Journal of Phycology 56(6): 1601–1613. 10.1111/jpy.13064

Kützing F. 1844. — Die kieselschaligen Bacillarien oder Diatomeen. Nordhausen, 152 p.

Lai G. G., Beauger A., Wetzel C. E., Padedda B. M., Voldoire O., Lugliè A., Allain E. & Ector L. 2019. — Diversity, ecology and distribution of benthic diatoms in thermo-mineral springs in Auvergne (France) and Sardinia (Italy). Peerj 7: e7238. 10.7717/peerj.7238

Lange-Bertalot H. & Krammer K. 1989. — *Achnanthes*, eine Monographie der Gattung mit Definition der Gattung *Cocconeis* und Nachträgen zu den Naviculaceae. Bibliotheca Diatomologica 18: 1–393.

Lange-Bertalot H. & Ruppel M. 1980. — A revision of some taxonomically most problematic groups in *Achnanthes* Bory, important from the ecological point of view. Algological Studies 26: 1–31.

Lange-Bertalot H., Külbs K., Lauser T., Nörpel-Schempp M. & Willmann M. 1996. — Diatom taxa introduced by Georg Krasske: Documentation and revision. Iconographia Diatomologica 3: 1–358.

Letunic I. & Bork P. 2024. — Interactive Tree of Life (iTOL) v6: recent updates to the phylogenetic tree display and annotation tool. Nucleic Acids Research 52 (W1): W78–W82. 10.1093/nar/gkae268

Liu Q., Xiang Y., Yu P., Xie S. & Kociolek J. P. 2020. — New and interesting diatoms from Tibet. II. Description of two new species of monoraphid diatoms. Diatom Research 35 (4): 353–361. 10.1080/0269249X.2020.1810781

Mann D. G. 1999. — The species concept in diatoms. Phycologia, 38(6): 437–495. 10.2216/i0031-8884-38-6-437.1

Mann D. G. 2010. — Discovering diatom species: is a long history of disagreements about species-level taxonomy now at an end? Plant Ecology and Evolution 143 (3): 251–264. 10.5091/plecevo.2010.405

Meinzer O. E. 1923. — Outline of Ground-Water Hydrology, with Definitions: Us Government Printing Office, Washington D.C. 71 p.

Minh B. Q., Nguyen M. A. T. & Von Haeseler A. 2013. — Ultrafast Approximation for Phylogenetic Bootstrap, Molecular Biology and Evolution 30(5): 1188–1195. 10.1093/molbev/mst024

Na X., Liu J., Zhang Y., Kociolek J. P., Kulikovskiy M., Lu X., Sui F., Zhu H., Liu G. & Fan Y. 2024. — A new species of genus *Crenotia* (Bacillariophyta) from Tibet, China. PhytoKeys 237: 23. 10.3897/phytokeys.237.112939

Nakov T., Beaulieu J. M. & Alverson A. J. 2018. — Accelerated diversification is related to life history and locomotion in a hyperdiverse lineage of microbial eukaryotes (Diatoms, Bacillariophyta). New Phytologist 219 (1): 462–473. 10.1111/nph.15137

Nawrocki E. P. 2009. — Structural Rna Homology Search and Alignment using Covariance Models. Phd Thesis, Washington University in St. Louis., 256 p. https://openscholarship.wustl.edu/etd/2561-206

Poretzky V. S. & Anisimova N. V. 1933. — *Materialy k ekologii diatomovykh Starorusskikh solenykh vodoemov* [Materials on Ecology of the Diatoms of Saltwater Basins of Staraia Russa]. Issledovaniia ozer Soyuza S.S.R. Gosudarstvennyi Gidrologicheskii Institut Leningrad 2: 31–66.

R Core Team 2021. — R: A language and environment for statistical computing. R Foundation for Statistical Computing Vienna, Austria.

Rabenhorst L. 1853. — Die Süsswasser-Diatomaceen (Bacillarien.) Für Freunde der Mikroskopie. Eduard Kummer, Leipzig, 1–72.

Rabenhorst L. 1864. — Flora Europaea Algarum Aquae Dulcis et Submarinae. Sectio I. Algas Diatomaceas Complectens, cum Figuris Generum Omnium Xylographice Impressis. Apud Eduardum Kummerum, Lipsiae [Leipzig], 1–359.

Riaux-Gobin C., Saenz-Agudelo P., Górecka E., Witkowski A., Daniszewska-Kowalczyk G. & Ector L. 2021. — *Cocconeis vaiamanuensis* sp. nov. (Bacillariophyceae) from Raivavae (South Pacific) and allied taxa: Ultrastructural specificities and remarks about the polyphyletic genus *Cocconeis* Ehrenberg. Marine Biodiversity 51 (2): 29. 10.1007/s12526-020-01154-9

Rioual P., Ector L. & Wetzel C. E. 2019. — Transfer of *Achnanthes hedinii* Hustedt to the genus *Crenotia* Wojtal (Achnanthidiaceae, Bacillariophyceae). Notulae Algarum 106: 1–6.

Round F. E., Crawford R. M. & Mann D. G. 1990. — The Diatoms. Biology and Morphology of the Genera. Cambridge University Press, Cambridge, 1–747 p.

Shimodaira H. 2002. — An approximately unbiased test of phylogenetic tree selection. Systematic Biology 51(3): 492–508. 10.1080/10635150290069913

Simonsen R. 1987. — Atlas and Catalogue of the Diatom Types of Friedrich Hustedt. J. Cramer in der Gebrüder Borntraeger Velagsbuchhandlung., Berlin & Stuttgart, 525 p.

Smith M. R. 2022. — Using information theory to detect rogue taxa and improve consensus trees. Systematic Biology 71(5): 1088–1094. 10.1093/sysbio/syab099

Taylor J., Harding W. & Archibald C. 2007. — A methods manual for the collection, preparation and analysis of diatom samples. Version 1: 60.

Thomas E. W., Stepanek J. G. & Kociolek J. P. 2016. — Historical and current perspectives on the systematics of the ‘enigmatic’diatom genus *Rhoicosphenia* (Bacillariophyta), with single and multi-molecular marker and morphological analyses and discussion on the monophyly of ‘monoraphid’diatoms. PLos One 11(4): e0152797. 10.1371/journal.pone.0152797

Thüs H., Muggia L., Pérez-Ortega S., Favero-Longo S. E., Joneson S., O’Brien H., Nelsen M. P., Duque-Thüs R., Grube M., Friedl T., Brodie J., Andrew C. J., Lücking R., Lutzoni F. & Gueidan C. 2011. — Revisiting photobiont diversity in the lichen family Verrucariaceae (Ascomycota). European Journal of Phycology 46 (4): 399–415. 10.1080/09670262.2011.629788

Trifinopoulos J., Nguyen L.-T., Von Haeseler A. & Minh B. Q. 2016. — W-IQ-TREE: a fast online phylogenetic tool for maximum likelihood analysis. Nucleic Acids Research 44 (W1): W232–W235. 10.1093/nar/gkw256

Trobajo R. & Mann D. G. 2019. — A rapid cleaning method for diatoms. Diatom Research 34 (2): 115–124. 10.1080/0269249X.2019.1637785

Urbánková P. & Veselá J. 2013. — DNA-barcoding: A case study in the diatom genus *Frustulia* (Bacillariophyceae). *Nova Hedwigia*, Beiheft 142 (3-4): 147–162.

Vences M., Patmanidis S., Kharchev V. & Renner S. S. 2022. — Concatenator, a user-friendly program to concatenate Dna sequences, implementing graphical user interfaces for Mafft and FastTree. Bioinformatics Advances 2 (1): vbac050. 10.1093/bioadv/vbac050

Veselá J., Urbánková P., Černá K. & Neustupa J. 2012. — Ecological variation within traditional diatom morphospecies: diversity of *Frustulia rhomboides* sensu lato (Bacillariophyceae) in European freshwater habitats. Phycologia 51 (5): 552–561. 10.2216/11-101.1

Wilm A., Higgins D. G. & Notredame C. 2008. — R-Coffee: a method for multiple alignment of non-coding RNA. Nucleic Acids Research 36 (9): e52–e52. 10.1093/nar/gkn174

Wojtal A. Z. 2013. — *Bibliotheca Diatomologica, Band 59*, Species composition and distribution of diatom assemblages in spring waters from various geological formations in southern Poland, 436 p.

Yoon H. S., Hackett J. D. & Bhattacharya D. 2002. — A single origin of the peridinin- and fucoxanthin-containing plastids in dinoflagellates through tertiary endosymbiosis. Proceedings of the National Academy of Sciences of the United States of America 99 (18): 11724–11729. 10.1073/pnas.172234799

