## Supplementary figures and images for "Inferring evolutionary relationships among *Crenotia* species (Bacillariophyta): Evidence from natural populations and monoclonal strains from Slovakia"

### Fig. S1

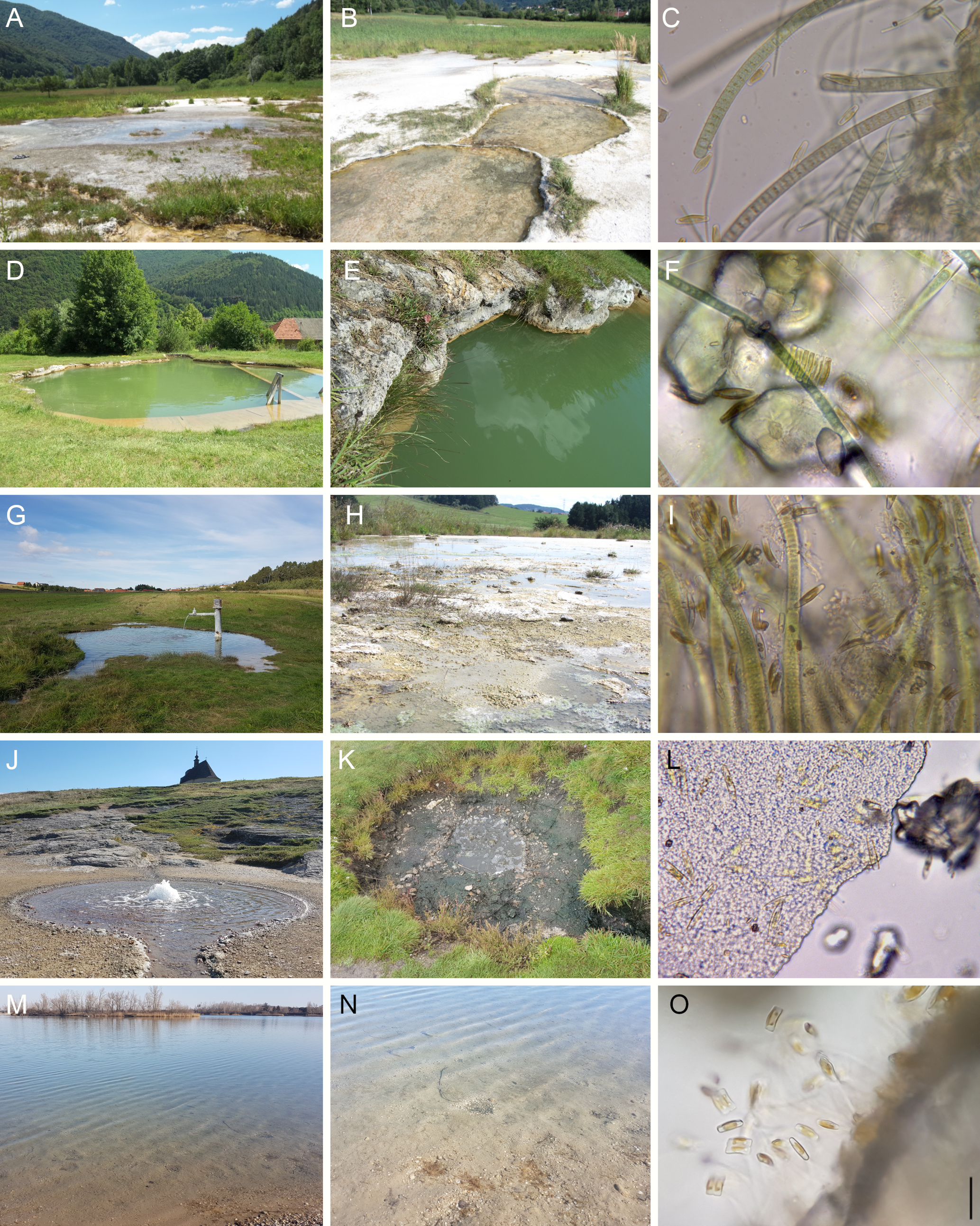

### Fig. S2

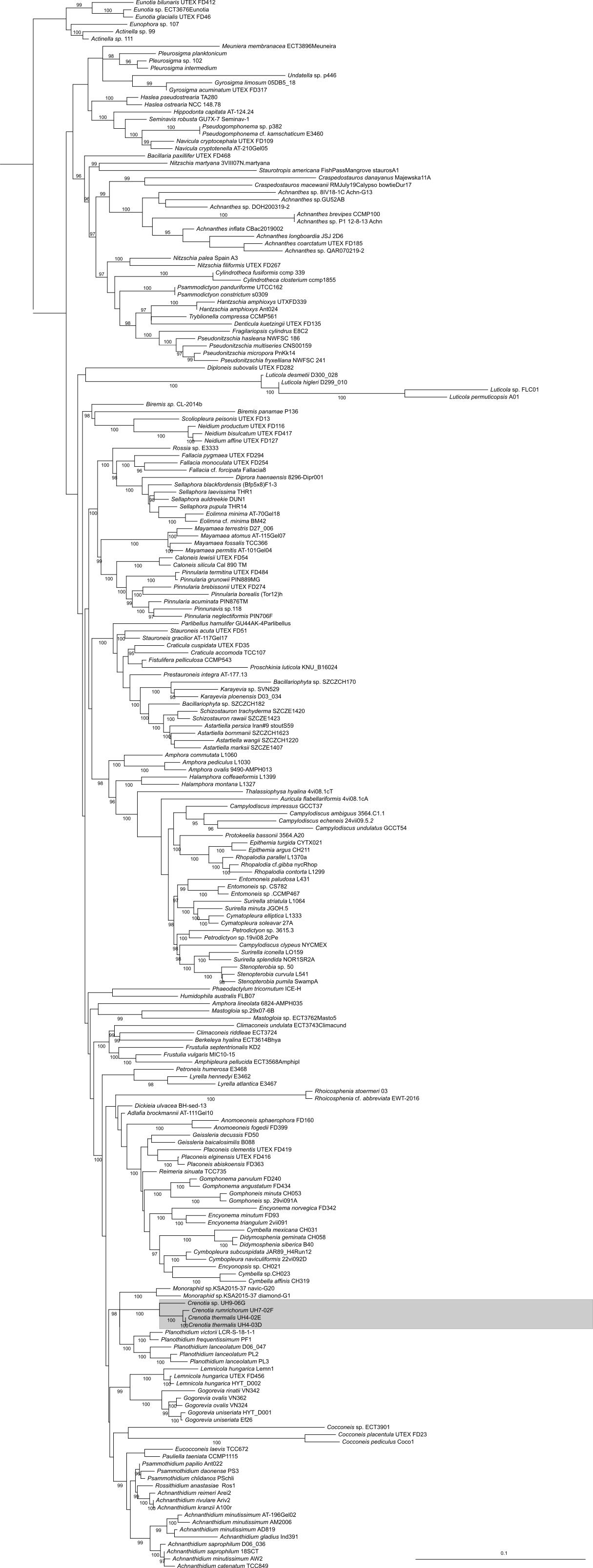
